# SuPreMo: a computational tool for streamlining *in silico* perturbation using sequence-based predictive models

**DOI:** 10.1101/2023.11.03.565556

**Authors:** Ketrin Gjoni, Katherine S. Pollard

**Affiliations:** Gladstone Institute of Data Science and Biotechnology, San Francisco, CA 94158, USA; Department of Epidemiology & Biostatistics, University of California, San Francisco, CA 94158, USA; Chan Zuckerberg Biohub, San Francisco, CA 94158, USA

## Abstract

Computationally editing genome sequences is a common bioinformatics task, but current approaches have limitations, such as incompatibility with structural variants, challenges in identifying responsible sequence perturbations, and the need for vcf file inputs and phased data. To address these bottlenecks, we present <u>S</u>equence M<u>u</u>tator for <u>Pre</u>dictive <u>Mo</u>dels (SuPreMo), a scalable and comprehensive tool for performing *in silico* mutagenesis. We then demonstrate how pairs of reference and perturbed sequences can be used with machine learning models to prioritize pathogenic variants or discover new functional sequences.

**Availability and Implementation:** SuPreMo was written in Python, and can be run using only one line of code to generate both sequences and 3D genome disruption scores. The codebase, instructions for installation and use, and tutorials are on the Github page: https://github.com/ketringjoni/SuPreMo/tree/main.

**Supplementary information:** Supplementary data are available at Bioinformatics online.

## 1 Introduction

Many machine learning (ML) models have been developed that predict cellular profiles from input DNA sequences (Supp Table 1). These sequence-to-profile models can predict biological features—including gene expression (Enformer (Avsec *et al*. 2021a), ExPecto (Zhou *et al*. 2018), Xpresso (Agarwal and Shendure 2020)), genome folding (Akita (Fudenberg, Kelley and Pollard 2020), C.origami (Tan *et al*. 2023), DeepC (Schwessinger *et al*. 2020), ORCA (Zhou 2022)), chromatin accessibility (Basenji (Kelley 2020), Basset (Kelley, Snoek and Rinn 2016)), and epigenetic marks (DeepFIGV (Hoffman *et al*. 2019), HyenaDNA (Nguyen *et al*. 2023), Sei (Chen *et al*. 2022a))—with incredible accuracy. These approaches are becoming increasingly popular for exploring biological questions at lower cost and higher throughput than experimental methods allow, and to address questions that are not possible to test experimentally. One exciting potential is to use sequence-to-profile models in tandem with *in silico* mutagenesis (ISM), in order to investigate how genomic alterations alter cellular profiles. This strategy generates testable, causal hypotheses about genotype-phenotype relationships (Chen *et al*. 2022b). ISM has been applied to the genomes of modern humans, archaic hominins (McArthur, Rinker and Capra 2021), and other species (Keough *et al*. 2023) to prioritize putative pathogenic variants for experimental studies (Benegas, Batra and Song 2023), decode the grammar of noncoding DNA sequences (Deng *et al*. 2023), discover new sequence motifs (Avsec *et al*. 2021b), design tissue-specific enhancers (Gosai *et al*. 2023), and uncover novel roles of sequence elements (Gunsalus, Keiser and Pollard 2023).

In theory, ISM is very high-throughput, making it feasible to quantify the effects of a large set of sequence perturbations, such as all variants in an individual’s genome or a cohort of patients. However, the application of ISM at scale is currently limited by the process of generating sequences with and without perturbations. Existing tools incorporate variants into a reference genome in ways that are not compatible with ISM (bcftools consensus (Li 2011), GATK FastaAlternateReferenceMaker (Van der Auwera and O’Connor 2020), perEditor (Rivas-Astroza *et al*. 2011), etc.). One of the biggest limitations is that they incorporate all variants from an input variant call format (vcf) (Danecek *et al*. 2011) file into a single output fasta file, making it very difficult to isolate the effects of individual variants. Workarounds, such as generating an independent vcf file for each variant (or variant combination) and looping over these or post-processing the output fasta file to include one variant per locus, are extremely inefficient. Secondly, existing tools are made largely for single nucleotide polymorphisms (SNPs) or small insertions or deletions (indels), and cannot accommodate symbolic alleles—annotations in vcf files for structural variants (SVs). A possible workaround is to convert symbolic alleles into sequences by extracting them from a reference genome, but this becomes infeasible with large structural variants due to limitations with both variant complexity and memory allocation. One tool (perEditor(Rivas-Astroza *et al*. 2011)) is compatible with some complex variants but is not comprehensive and has stringent requirements. Finally, existing tools require the perturbations to be in a vcf format, which means that pseudo input files must be generated if one wishes to apply ISM to custom or simulated sequences (e.g., deleting all motifs for a given transcription factor or creating synthetic enhancers).

Due to these limitations, it is common practice for ISM practitioners to write their own code to generate input sequences for ISM studies. Indeed, the codebases for several ML models include code examples or frameworks for performing ISM (Enformer, Sei (Chen *et al*. 2022a), Basset), but these are restricted to simple variants (SNPs and indels) and do not generate sequence files for input into other models. SVs make good candidates for ISM since they span larger regions and are more likely to be damaging to the genes, regulatory regions or active sites they overlap or neighbor. For example, noncoding SVs have been shown to lead to cancer and developmental disorders by disrupting genomic contacts of key genes (Paik, Maule and Gallo 2021). SVs also alter more base pairs of the genome than any other type of genetic variation (1000 Genomes Project Consortium *et al*. 2015). One major challenge with SVs is that, to adhere to the fixed length input requirements of most ML models, input sequences must be padded, and consequently, model outputs require un-padding and masking. Another consideration is that—due to both biological effects and model artifacts related to making predictions for fixed width genomic windows—models can be highly sensitive to small changes in the input, such as masking, padding, and variant position in the window. Therefore, it is important to make predictions for augmented input sequences (shifted and/or reverse complement sequences) and evaluate them consistently across perturbations. Thus, incorporating perturbations into a reference genome becomes increasingly complicated and error-prone as variants get larger and more complex.

## 2 Tool description

To address these challenges, we developed SuPreMo, a framework for generating perturbed sequences for input into predictive models that is scalable, flexible, and comprehensive (Fig 1A). SuPreMo, which incorporates variants into the human reference genome one at a time and generates model-ready sequences (Supp Fig 1A), was extended to SuPreMo-Akita, which inputs those sequences into Akita (Fudenberg, Kelley and Pollard 2020), an ML model that predicts chromatin contact maps, and generates scores that measure variants’ disruption to those maps (Supp Fig 1C). Both tools accept a variety of variant files—vcf (version 4.1 and 4.2), txt, bed-like, and tsv (generated from AnnotSV (Geoffroy *et al*. 2018))—making them flexible for use with real or synthetic perturbations. The following variant types (marked by their Manta (Chen *et al*. 2016) abbreviations) are supported: SNPs, indels, deletions (DEL), duplications (DUP), inversions (INV), and complex rearrangements with breakends (BNDs). Across a variety of datasets, including the 1K Genome Project, SuPreMo makes it possible to analyze over 50% of SVs that would not be accessible with existing tools (Fig 1B). In particular, symbolic alleles are now easily and uniformly processed (Fig 1B, orange). On the other hand, insertions, which make up <20% of SVs, remain inaccessible for sequence-based models since the precise inserted sequence is not provided by SV calling methods (Fig 1B, gray).

**Figure 1:**
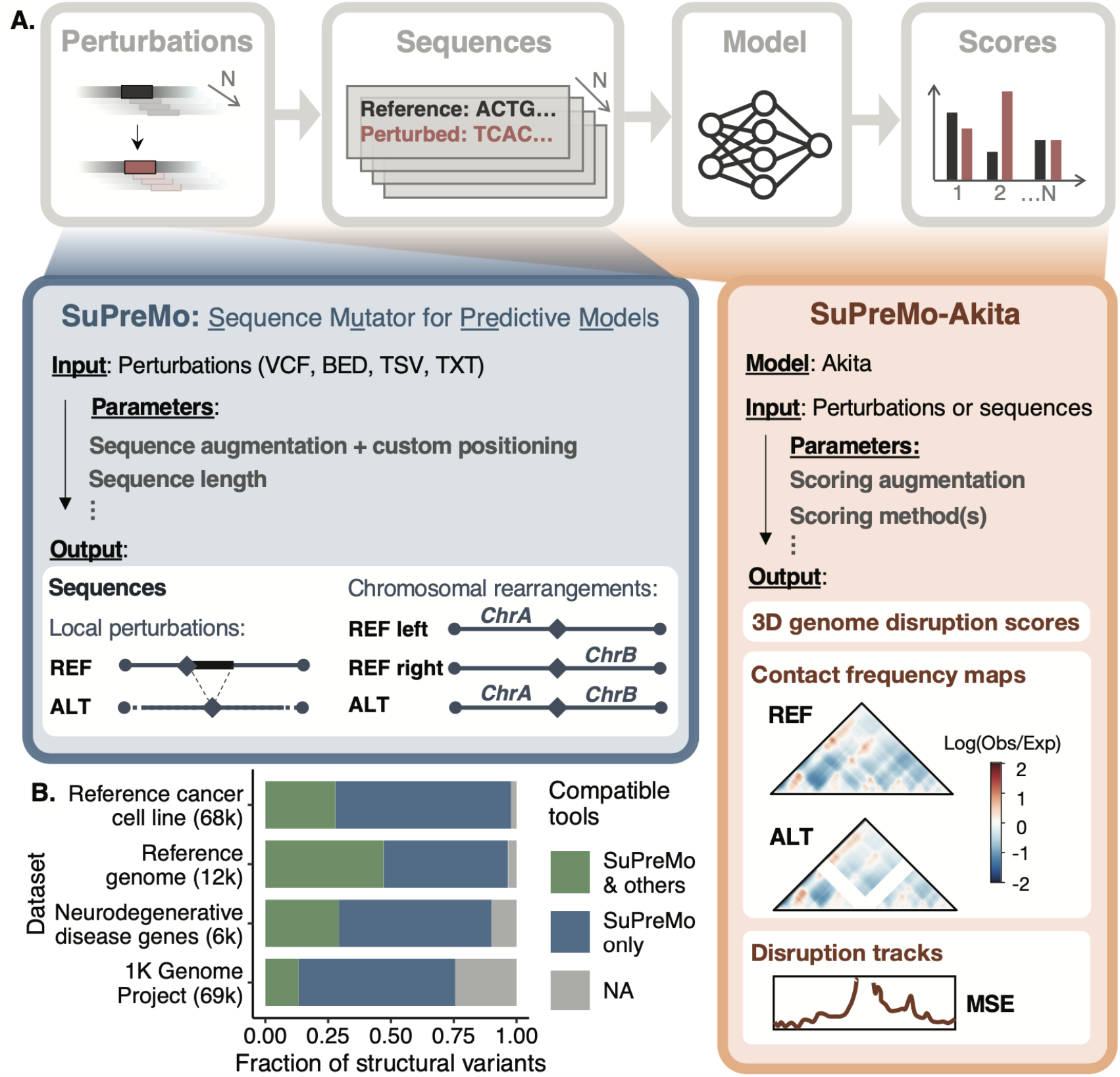
A: Schematic representation of SuPreMo. SuPreMo generates sequences by incorporating perturbations into the hg38 human reference genome. SuPreMo-Akita applies Akita to those sequences and generates 3D genome disruption scores (effect size of each perturbation) and, optionally, disruption tracks and predicted contact frequency maps. Parameters and outputs are specified. REF: derived from reference allele; ALT: derived from alternate allele; Log(Obs/Exp): log of observed over expected contacts; MSE: mean squared error. B: Categorization of SVs based on the ability of SuPreMo and other existing tools to incorporate them into a reference genome. SVs that other tools can already process include small indels (green); SVs that only SuPreMo can process include deletions, duplications, inversions, and chromosomal rearrangements (navy); SVs that no tool can process include insertions and copy number variants (CNVs) because the exact sequence is not provided by upstream variant calling pipelines (gray). Datasets are WGS/WES from healthy and disease individuals: a reference cancer cell line (Talsania *et al*. 2022), a reference genome (Zook *et al*. 2020), neurodegenerative disease gene sequences (Kaivola *et al*. 2023), and the 1K Genome Project (Mahmoud *et al*. 2019).

SuPreMo provides flexibility through the following parameters: prediction window shift, output sequence length, maximum variant length, and reverse complement. Variants are centered in the output sequences unless they are too close to chromosome arm ends (Supp Fig 1B) or the shift parameter is used, which moves the prediction window around the variant and therefore changes its position in the sequence (Supp Fig 2). The length of the generated sequences can be customized to match the required input of the model of interest (Supp Table 1). For SV inputs, the maximum SV length considered is limited to two thirds of the input sequence, unless otherwise specified by the user. Lastly, the reverse complement of the generated sequence can be outputted. Generated sequences are accompanied by the relative position of the variant in each sequence. Thus, SuPreMo is a flexible tool for performing ISM that can be applied across sequence-based ML models.

SuPreMo-Akita generates an array of 3D genome disruption scores, predicted contact frequency maps for reference (wild type) and alternate (perturbed) sequences, and genomic tracks of disruption scores across the prediction window. Akita predicts contact frequency maps for a ∼1 megabase (Mb) input sequence at a ∼2 kilobase (kb) resolution. SuPreMo-Akita inputs variants as described above and optionally also takes in already generated sequences. Since methods for scoring contact maps are biased and sometimes only target certain features, we have made available 13 different predefined metrics (Gunsalus *et al*. 2023) to use with this tool, with the defaults being the most common measures: mean squared error (MSE) and Spearman’s rank correlation coefficient (referred to here as just correlation). To assess the robustness of the generated disruption scores, the augmentation parameter optionally provides averages of scores from standard sequences, sequences with −1 bp and +1 bp shifts, and reverse complement sequences, or any other augmentations specified. Each generated map will be accompanied by the start genomic position and the relative bin that the variant lies in.

Lastly, we considered computational efficiency. To enable customization to different hardware, the user can choose the number of rows to be processed at a time from the input file and what outputs to request, keeping in mind storage and memory limitations. We measured the run time, peak memory, and size of outputs on 3 GHz CPUs using a set of 100-1000 SVs of different types from the reference cancer cell line in Figure 1B. SuPreMo-Akita is fast and easily scaled up—with the augmentation parameter it takes approximately 19 seconds and accumulates ∼ 0.15 megabytes (MB) of memory per variant (Supp Table 2).

We implemented SuPreMo using two models, although our framework is extendable to any model utilizing genome sequences as input. First, we used SuPreMo with DeepSEA (Zhou and Troyanskaya 2015) to rank a set of CTCF deletions based on their predicted effect on epigenetic marks. Second, we used SuPreMo-Akita on cancer SVs (Supp Fig 3A). SVs were scored using MSE and correlation, and the top 3 scoring variants for each SV type and scoring method were selected (Supp Fig 3B). We separately ranked variants by their type because their 3D genome disruption scores vary, and by the scoring method because each has unique biases. Using SuPreMo-Akita, contact frequency maps and disruption tracks were generated for these selected SVs and the most interesting variants, based on the structures they disrupt, were chosen (Supp Fig 3C). This method prioritized a deletion of an insulated site that is predicted to cause increased contact frequency between neighboring regions (Supp Fig 3C, left panel). Step-by-step instructions for both implementations are available on Github.

## 3 Conclusion

SuPreMo is a software tool that facilitates ISM with predictive models and extends this principle with Akita to predict scores for 3D genome folding disruption. Potential use cases include scoring all variants in an individual or cohort for disruption to genome folding, generating predicted contact frequency maps to explore the effects of noncoding variants on regulatory interactions, performing ISM to evaluate or discover sequence motifs using Akita, and, more broadly, generating sequences for input into predictive models of interest to evaluate variant effects. SuPreMo is scalable to a large number of variants and only limited by the storage capacity the user has for the expected outputs. Overall, SuPreMo allows for easy, fast and broadly applicable analysis of simple variants, SVs, and chromosomal rearrangements in the context of sequence-based predictive models.

## Acknowledgements

We thank Shu Zhang for helpful discussion and feedback and for reviewing the manuscript. We thank Maureen Pittman, Laura Gunsalus, and Shuzhen Kuang for helpful insight on using Akita.

