## Supplemental Table 1 for "SuPreMo: a computational tool for streamlining *in silico* perturbation using sequence-based predictive models"

### Supplementary Table 1

**Sequence-based predictive models and their specifications**

| **#** | **Model** | **Input sequence length (bp)** | **Predicted output** | **Output format** | **PMID (or DOI^b^)** |
| --- | --- | --- | --- | --- | --- |
| 1 | Akita | 1,048,576 | Contact frequency map | Matrix | 33046897 |
| 2 | Basenji | 131,072 | Various chromatin profiles (CAGE, ATACseq, ChIPseq) | Genomic tracks | 32687525 |
| 3 | Basset | 600 | DNA accessibility | Genomic tracks | 27197224 |
| 4 | Borzoi | 524 | Various chromatin profiles (CAGE, RNA-seq, DNase or ChIP-seq) | Genomic tracks | 08.30.555582 |
| 5 | BPnet | 1000 | Various chromatin profiles (TF ChIP-nexus) | Genomic tracks | 33603233 |
| 6 | C.Origami^a^ | 2,097,152 | Contact frequency map | Matrix | 36624151 |
| 7 | DanQ | 1000 | Various chromatin profiles (DNase, ChIPseq) | Genomic tracks | 27084946 |
| 8 | DeepC | 1,005,000 | Contact frequency map | Matrix | 33046896 |
| 9 | DeepFIGV | 300-2000 | Various chromatin profiles (DNase, histone ChIPseq) | Genomic tracks | 31544924 |
| 10 | DeepSea | 1000 | Various chromatin profiles (TFBS, DNase, histone ChIPseq) | Values | 26301843 |
| 11 | DeepTACT^a^ | 5,000-20,000 | Contact frequency (only for regulatory elements) | Values | 30869141 |
| 12 | Enformer | 196,608 | Various chromatin profiles (CAGE, DNase, ChIPseq) | Genomic tracks | 34608324 |
| 13 | ExPecto | 40,000 | Gene expression | Values (log RPKM) | 30013180 |
| 14 | ExPectoSC | 40,000 | Gene expression | Values (log RPKM) | 37703883 |
| 15 | HyenaDNA | 1,000-8,000 | Various chromatin profiles (TFBS, DHS, histone ChIPseq at central 200 bp) | Values | 37426456 |
| 16 | Malinois | 770,000 | Gene expression (based on CRE activity) | Values (sequence contribution scores) | 37609287 |
| 17 | ORCA | 1,000,000- 256,000,000 | Contact frequency map | Matrix | 35551308 |
| 18 | Puffin-D | 100,000 | Gene expression (transcription initiation signals) | Genomic tracks | 06.27.546584 |
| 19 | Sei | 4,000 | Various chromatin profiles (x21,907) | Sequence classes | 35817977 |
| 20 | Seq-GraphReg | 6,000,000 | Gene expression | Genomic tracks | 35396274 |
| 21 | seq2cells | 200,000 | Gene expression | Values | 07.26.550634 |
| 22 | TREDnet | 2,000 | Various chromatin profiles (TFBS, DNase, histone ChIPseq) | Values | 37603758 |
| 23 | Xpresso | 10,500 | Gene expression | Genomic tracks | 32433972 |

^a^require additional input(s)

^b^DOIs start with 10.1101/2023.
