## Supplemental Table 2 for "SuPreMo: a computational tool for streamlining *in silico* perturbation using sequence-based predictive models"

**Supplementary Table 2**

**SuPreMo and SuPreMo-Akita specifications**

|  | **SuPreMo** | **SuPreMo-Akita** | **SuPreMo-Akita** |
| --- | --- | --- | --- |
| **Parameters** | --seq_len 10000  --shift -1 0 1  --revcomp add_revcomp | --augment | --augment  --get_maps  --get_tracks |
| **Number of variants** | 1000 | 100 | 100 |
| **Time (s)^a^** | 12 +/- 0.4 | 1,904 +/- 29.8 | 2,313 +/- 48.0 |
| **Peak memory (KB)^a^** | 116,805 +/- 0.4 | 526,765 +/- 7,900 | 1,772,261 +/- 4,126 |
| **Size of output (KB)** | Sequences: 83,014 | Scores: 9.5 | Maps: 638,029  Tracks: 2,935 |

^a^Mean +/- standard deviation across 30 runs.
