## Supplemental Figure 1 for "SuPreMo: a computational tool for streamlining *in silico* perturbation using sequence-based predictive models"

### Supplementary Figure 1

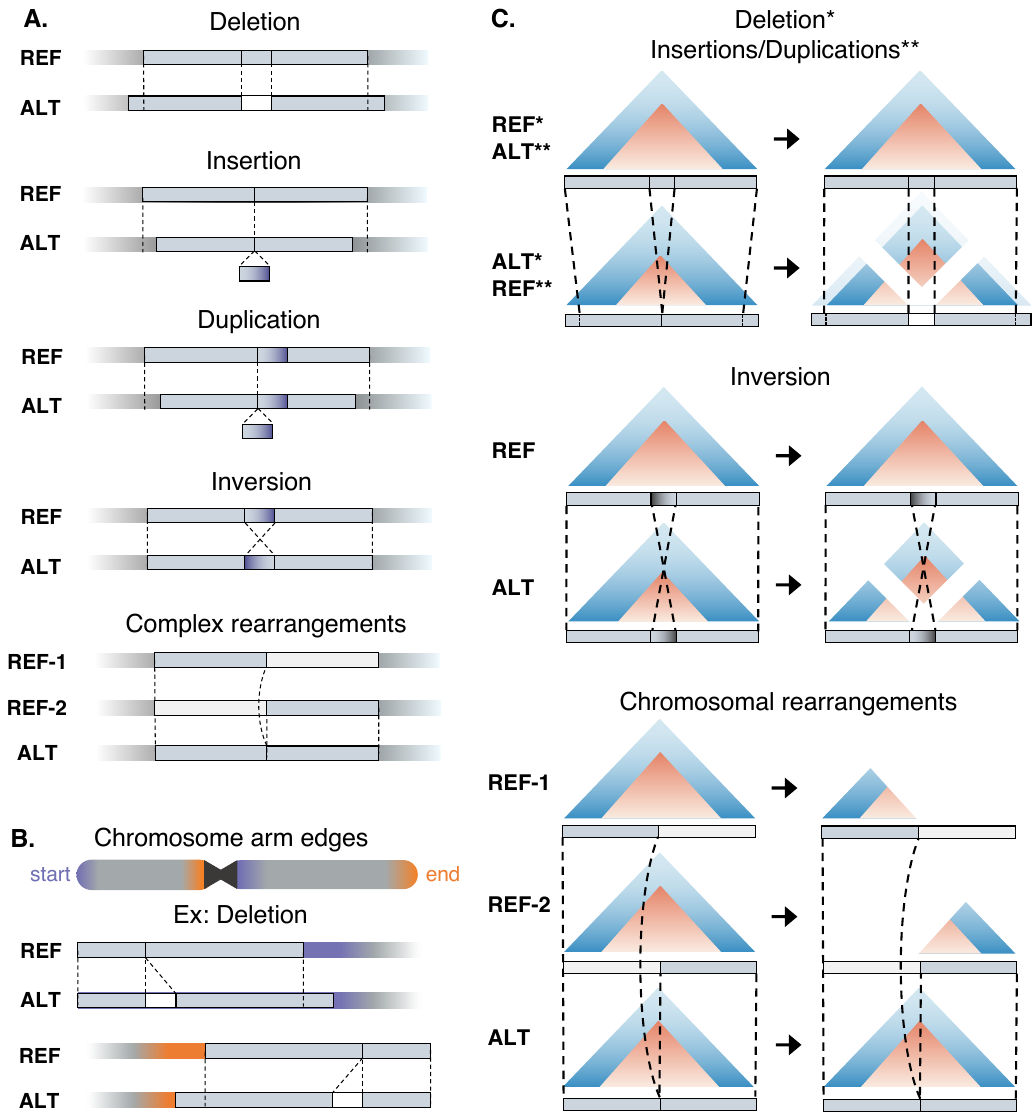

**Schematic of getting sequences and maps.** A: Schematic of how SuPreMo incorporates variants into a reference genome. Given a variant, SuPreMo generates reference sequence(s) (REF) and an alternate, perturbed sequence (ALT). B: Schematic of how SuPreMo-Akita adjusts contact frequency maps before comparing them to generate a perturbation score. Examples are categorized by variant type. Left panel contains maps for each allele as predicted by Akita, and the right panel shows those maps processed and ready to compare. Gray: reference genome; light blue boxes: sequences generated; white boxes: sequence deleted; dotted lines: matching positions in REF and ALT; white areas in maps: padded regions or regions with no prediction; faded areas in maps: cropped regions.
