## Supplemental Figure 2 for "SuPreMo: a computational tool for streamlining *in silico* perturbation using sequence-based predictive models"

### Supplementary Figure 2

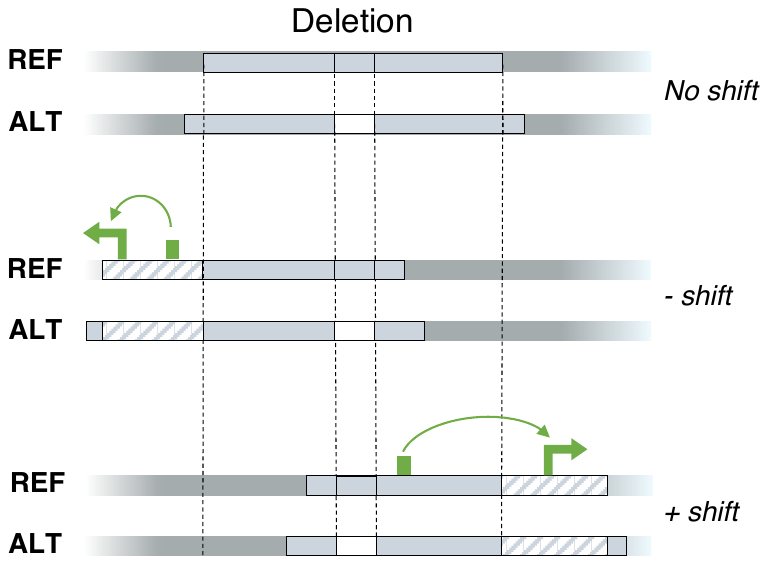

### Schematic of the shift parameter. Variants are by default centered in the generated sequences, as the deletion shown on the top. The shift parameter can be used to shift the prediction window upstream (negative shift, middle) or downstream (positive shift, bottom). A potential incentive for this is to include certain genes or regulatory elements that otherwise would not have been included in the window.
