## Supplemental Figure 3 for "SuPreMo: a computational tool for streamlining *in silico* perturbation using sequence-based predictive models"

### Supplementary Figure 3

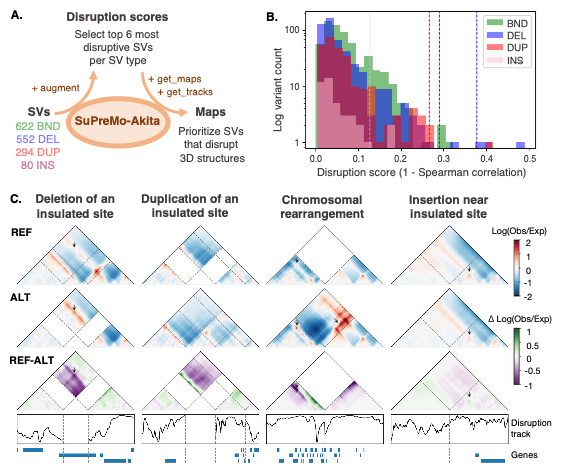

**Example application of SuPreMo-Akita on tumor SVs.** A: Schematic of scoring SVs downloaded from Talsania et al. 2022 with SuPreMo-Akita to get scores and then scoring a subset to get maps and tracks. B: Distribution of disruption scores (MSE also calculated but not shown) for SVs in A. Dashed line marks the threshold of the top 3 scoring variants for each type. C: Contact frequency maps and disruption tracks generated by SuPreMo-Akita for one selected, highly disruptive variant for each SV type. White areas in maps: padded regions or regions with no prediction; arrows: regions with changed contact.
